# A Van Gogh/Vangl tyrosine phosphorylation switch regulates its interaction with core Planar Cell Polarity factors Prickle and Dishevelled

**DOI:** 10.1101/2022.05.25.493488

**Authors:** Ashley C. Humphries, C. Clayton Hazelett, Claudia Molina-Pelayo, Danelle Devenport, Marek Mlodzik

## Abstract

Epithelial tissues can be polarized along two axes, in addition to apical-basal polarity they are often also polarized within the plane of the epithelium, known as planar cell polarity (PCP). PCP depends upon the conserved Wnt/Frizzled (Fz) signaling factors, including Fz itself and Van Gogh (Vang/Vangl). Here, taking advantage of the complementary features of *Drosophila* wing and mouse skin PCP establishment, we dissect how Vang phosphorylation on a specific conserved tyrosine residue affects its interaction with two cytoplasmic core PCP factors, Dsh/Dvl and Pk. We demonstrate that Pk and Dsh/Dvl bind to Vang/Vangl in an overlapping region centered around this tyrosine. Strikingly, Vang/Vangl2 phosphorylation promotes its binding to Pk, a key effector of the Vang/Vangl complex, and inhibits its interaction with Dsh/Dvl, and thus phosphorylation of this tyrosine appears to promote the formation of the mature and stable Vang/Vangl-Pk complex during PCP establishment. Interestingly, as our single point mutations allow selective binding inhibition of either Dsh or Pk, we can demonstrate for the first time that Dsh interaction with Vang is physiologically important, as all single point mutations fail to rescue the *Vang* null mutant wing phenotype.

## Introduction

Cellular polarization is critical for the morphogenesis and function of organs and most tissues during development, with perturbation of cellular polarity and tissue organization implicated in numerous diseases. In particular, epithelial cells can be polarized in two axes, the ubiquitous epithelial polarity in the apical-basal axis and polarity orthogonal to the plane of the epithelium, with the latter being generally referred to as planar cell polarity (PCP) (e.g. reviewed in (Adler, 2012; Axelrod, 2020; Butler and Wallingford, 2017; Devenport, 2014; Goodrich and Strutt, 2011; Humphries and Mlodzik, 2018; Yang and Mlodzik, 2015)). Although PCP was initially discovered in epithelia, the cellular mechanisms controlling PCP are detected in many other cell types, including migratory mesenchymal cells or migratory neurons (reviewed in (Davey and Moens, 2017; Devenport, 2014; Koca et al., 2022). PCP establishment is mainly governed by members of the conserved Wnt/Frizzled-PCP pathway, often referred to as the Core PCP pathway (reviewed in (Adler, 2012; Axelrod, 2020; Butler and Wallingford, 2017; Devenport, 2014; Goodrich and Strutt, 2011; Humphries and Mlodzik, 2018; Yang and Mlodzik, 2015))

The core PCP factors, all originally discovered in *Drosophila*, include the atypical seven-pass transmembrane (TM) cadherin Flamingo (Fmi; Celsr in mammals), the seven-pass TM protein Frizzled (Fz; Fzd in vertebrates with several family members), and the four-pass trans-membrane protein Vang (Vangl1 and Vangl2 in mammals; a.k.a. *strabismus/stbm* in Drosophila and Xenopus). Besides these TM-proteins, the core PCP factors also include the three cytoplasmic proteins Dishevelled (Dsh; Dvl in mammals), Diego (Dgo; Inversin/Diversin in vertebrates), and Prickle (Pk) (e.g. reviewed in (Adler, 2012; Axelrod, 2020; Butler and Wallingford, 2017; Devenport, 2014; Goodrich and Strutt, 2011; Humphries and Mlodzik, 2018; Yang and Mlodzik, 2015)). The resulting initial outcome of the PCP signaling pathway is largely synonymous with asymmetric localizations of its core members. These become enriched into two complexes on opposite sides of a given cell, or in non-overlapping domains in migrating cells. In epithelial cells they generate an intracellular bridge to convey polarity across cell membranes from cell to cell across the tissue (see reviews above). As such the resulting complexes antagonize each other intra-cellularly and stabilize each other across membranes inter-cellularly. The resulting final asymmetric localization is generated through a dynamic process, including unstable intermediary subcomplexes. The ‘terminal’ asymmetric core PCP complex localization then directs spatially restricted downstream signaling events through cell-type specific effectors, leading to - depending on the tissue - cytoskeletal rearrangement, centriole/centrosome/ciliary positioning, migratory regulation, and/or nuclear read-outs (reviewed in (Adler, 2012; Axelrod, 2020; Butler and Wallingford, 2017; Carvajal-Gonzalez et al., 2016; Davey and Moens, 2017; Devenport, 2014; Humphries and Mlodzik, 2018; Koca et al., 2022; Yang and Mlodzik, 2015)).

Although the genetics of PCP have been discovered and initially functionally dissected in *Drosophila*, and are still best studied in flies (Adler, 2012; Goodrich and Strutt, 2011; Humphries and Mlodzik, 2018; Seifert and Mlodzik, 2007), the importance of PCP during vertebrate development and human disease has become widely recognized (reviewed in (Axelrod, 2020; Butler and Wallingford, 2017; Carvajal-Gonzalez et al., 2016; Devenport, 2014; Simons and Mlodzik, 2008; Wang and Nathans, 2007; Yang and Mlodzik, 2015). The individual PCP pathway components and their molecular interactions, leading to cellular polarization are highly conserved across metazoans up to mammals, with many different tissues displaying distinct PCP readouts or associated defects upon misregulation (see above reviews). For example, PCP directs polarized ciliary beating to generate fluid flow in the embryonic node, trachea, oviduct and brain ventricles. Failure to generate coordinated fluid flow in PCP mutant mice leads to left-right patterning defects, defective mucociliary clearance, sterility and hydrocephalus (Axelrod, 2020; Boutin et al., 2014; Ohata et al., 2014; Shi et al., 2014; Shinohara and Hamada, 2017; Tissir et al., 2010; Vladar et al., 2012; Wu and Mlodzik, 2017). Core PCP genes are essential for neural tube closure in mammals and pathological variants in PCP genes are strongly associated with neural tube defects in humans (Cai and Shi, 2014; Nikolopoulou et al., 2017; Wang et al., 2019). A notable example of PCP in mammals is the uniform alignment of body hairs across the skin surface, where the core PCP proteins direct polarized morphogenesis and tissue-wide alignment of hair follicles (Cetera et al., 2018; Devenport and Fuchs, 2008; Guo et al., 2004; Ravni et al., 2009).

Asymmetric distribution of PCP complexes is a direct consequence of their interactions during polarity establishment. It is observable in several tissues, ranging from cells of the *Drosophila* wing, where these complexes align to the proximal-distal axis, to for example mouse skin, where they align in the antero-posterior axis (Bastock et al., 2003; Das et al., 2004; Devenport and Fuchs, 2008; Strutt, 2001), also reviewed in (Adler, 2012; Devenport, 2014; Goodrich and Strutt, 2011; Humphries and Mlodzik, 2018; Seifert and Mlodzik, 2007)). Molecular interactions promote the formation of stable complexes at proximal (*Drosophila* wing) or anterior (mouse skin) cell membranes (Fmi/Celsr1-Vang/Vangl-Pk) and the equivalent distal or posterior cell surfaces (Fmi/Celsr1-Fz/Fzd-Dsh/Dvl), with Fmi (Celsr1 in mammals) forming a homotypic interaction across cells, which stabilizes these complexes intercellularly. Simple epithelia like the *Drosophila* wing display not only such highly coordinated PCP complex localization logic, but also an obvious and simple PCP read-out, with the formation of a single-actin based hair at the distal vertex of each cell pointing distally. Disruption of PCP establishment in the wing is easily observable by misorientation of the cellular hairs within the field of cells, or the formation of multiple cellular hairs in a single cell (rev in Adler, 2012; Goodrich and Strutt, 2011; Humphries and Mlodzik, 2018; Seifert and Mlodzik, 2007).

Among the core PCP factors the Vang/Vangl family proteins occupy a unique role, as they have been shown to physically interact with all other 5 core PCP factors (rev in (Adler, 2012; Goodrich and Strutt, 2011; Humphries and Mlodzik, 2018; Seifert and Mlodzik, 2007)) and also the A/B-polarity protein Scribble (Scrib) (Courbard et al., 2009; Montcouquiol et al., 2003). For example, all three cytoplasmic core factors, Dsh, Pk, and Dgo (and their respective vertebrate orthologues) have been shown to bind to the cytoplasmic C-terminal tail of Vang (e.g. see (Bastock et al., 2003; Jenny et al., 2005)). Similarly, Vang/Vangl proteins have been shown to associate with the trans-membrane factors Fmi/Celsr in cis and Fz in trans (Chen et al., 2008; Devenport and Fuchs, 2008; Stahley et al., 2021; Wu and Mlodzik, 2008). *Drosophila Vang* (a.k.a. *strabismus*/*stbm* in flies) has been identified by its strong PCP loss-of-function defects in wing and eye screens, and also demonstrated to cause a domineering non-autonomous phenotype affecting nearby wild-type cells through propagation of aberrant polarity from mutant cells outward (Taylor et al., 1998; Wolff and Rubin, 1998). Vang/Vangl genes encode four-pass transmembrane proteins with intracellular cytoplasmic amino- and carboxy-terminal regions. All cytoplasmic interactions have thus far been physically mapped to its C-terminal tail. It requires Pk and Scrib interactions for the formation of its stable complex, and can also interact with Dsh and Dgo, although these latter interactions are thought to be more transient and antagonistic to stable complex formation (Bastock et al., 2003; Courbard et al., 2009; Das et al., 2004; Jenny et al., 2005; Montcouquiol et al., 2003). Its mammalian homologs (*Vangl1 and Vangl2*, which is called *trilobite* in zebrafish) regulate all PCP processes studied in higher organisms (see for example (Jessen et al., 2002; Marlow et al., 1998; Montcouquiol et al., 2003), reviewed in (Wallingford, 2006; Ybot-Gonzalez et al., 2007)). Lethal mutations of both *Vangl1* and *Vangl2* have also been identified in human patients affected with *spina bifida* and *craniorachischisis* (Lei et al., 2010) and subsequently shown to affect PCP signaling and establishment (Humphries et al., 2020).

In order to better define the mechanisms underlying Vang/Vangl regulation and its interactions with downstream PCP factors, we investigated how phosphorylation of Vang/Vangl might affect and regulate its function. It was previously shown that Vang/Vangl proteins are serine/threonine phosphorylated in its N-terminal tail, which affects complex formation and stability (Chuykin et al., 2021; Gao et al., 2011; Kelly et al., 2016). However, it remains unclear how tyrosine phosphorylation might affect Vang/Vangl function and whether phosphorylation might regulate its interaction(s) with the cytoplasmic PCP factors. Here we describe a concerted effort using the *Drosophila* wing and mouse skin models to better define potential Vang/Vangl interactions as regulated by phosphorylation. We focus on a conserved phosphorylated C-tail tyrosine, which also resides within the broad region of Pk and Dsh/Dvl binding to Vang. Strikingly, the defined phosphorylation site resides in the newly defined overlapping binding regions of Pk and Dsh, and it regulates Vang/Vangl interactions with these core factors. Pk binds preferentially to the phosphorylated state, and Dsh to the unphosphorylated region. Our functional *in vivo* rescue follow-up studies demonstrate that binding of Vang to both effectors is physiologically relevant, as all single point mutations, which allow selective binding to either Pk or Dsh fail to rescue the *Vang* null mutant phenotype. With our defined single point mutation in Vang, we have shown the physiological relevance of the Dsh interaction with Vang for the first time. And further, we have identified means of regulating binding to antagonistic effectors during PCP establishment via a phosphorylation switch at a conserved site..

## Results

### Vang/Vangl2 proteins display a multitude of phosphorylations *in vivo*

Several studies have demonstrated that *Drosophila* Vang and mouse Vangl2 are phosphorylated on serine (S) and threonine (T) residues, and a functionally important S/T phosphorylation cluster has been defined (Chuykin et al., 2021; Gao et al., 2011; Kelly et al., 2016; Strutt et al., 2019). In addition, phosphorylation on a specific tyrosine (Y) has been shown to be important for correct Vangl2 trafficking in mammalian cells and *Drosophila* Vang *in vivo*, respectively (Guo et al., 2013; Kelly et al., 2016).

Our initial analyses performed in the Kelly et al. (2016) study have strongly suggested that additional Y residues are also likely to be phosphorylated. To better define the role of tyrosine phosphorylation in the context of Vang/Vangl function, we first verified that in *Drosophila* Vang is tyrosine phosphorylated *in vivo* (Figure 1A). Immunoprecipitation of Vang-Flagx3 from larval wing discs, showed a positive phospho-tyrosine signal, which importantly was reduced upon phosphatase treatment (Figure 1A; note that a down-shift in the mobility of Vang consistent with successful phosphatase treatment was observed). Second, importantly, Vang remained Y-phosphorylated in the Vang-Y341F mutant (Figure 1B). As this site has been defined in the past as phosphorylated, with Y341 (Y280/281 in *Vangl2*) promoting correct membrane trafficking (Guo et al., 2013; Kelly et al., 2016), these data suggested that other tyrosines must be phosphorylated in Vang/Vangl2 proteins *in vivo*.

**Figure 1.**
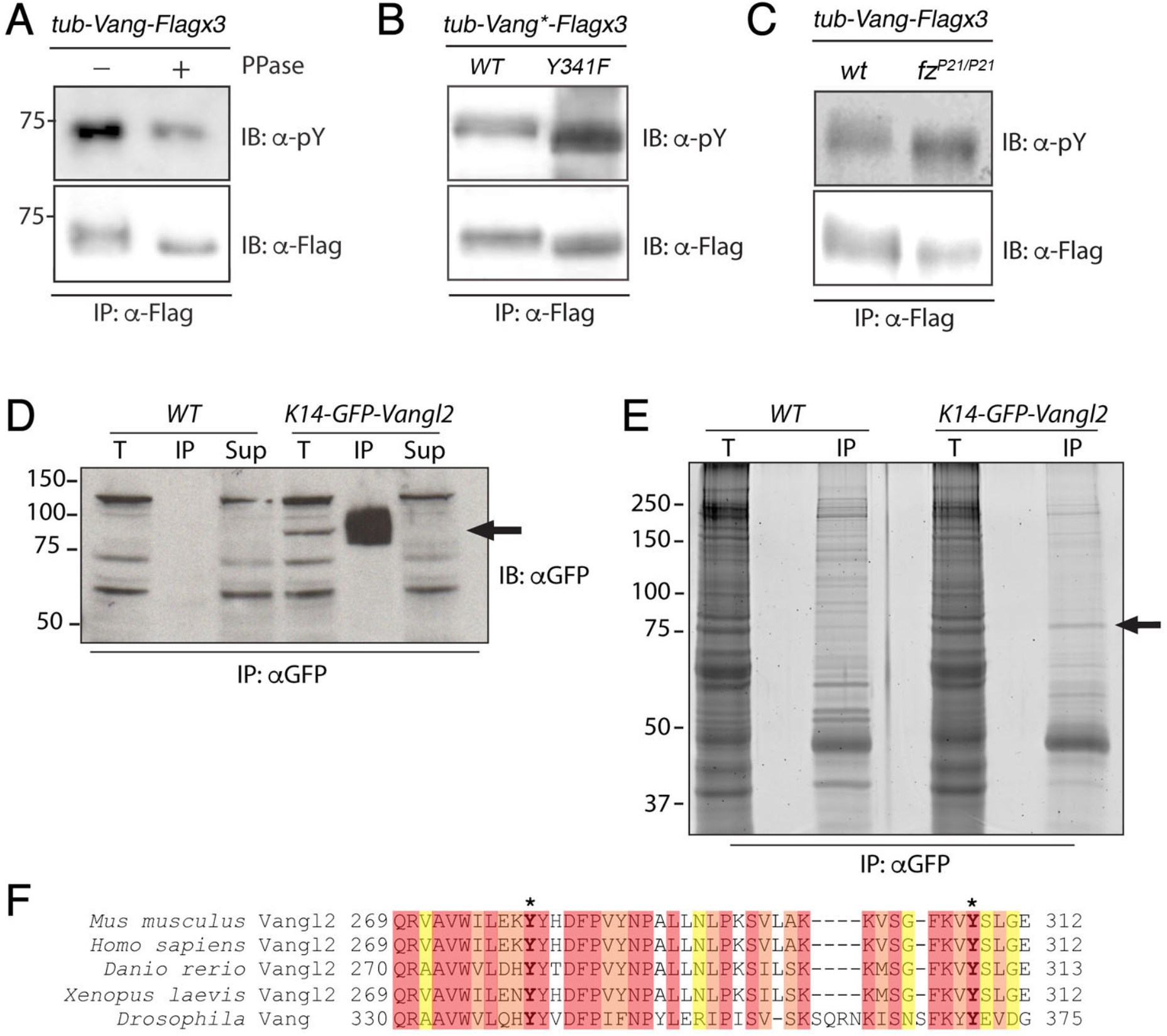
Vang is tyrosine phosphorylated in vivo. (**A-C**) Western blots of lysates from pupal wing discs showing that Vang is tyrosine phosphorylated *in vivo*. Note the reduction in signal of anti-pY with PPase treatment (**A**, upper panel) and the corresponding loss of a band shift (lower panel). (**B**) Tyrosine phosphorylation is maintained in the Vang-Y341F protein, suggesting other tyrosines to be phosphorylated; and tyrosine phosphorylation is independent of Fz, as anti-pY staining is not affected in homozygous mutants *fz*^*P21*^ null animals (**C**). **(D)** Immunoprecipitation of GFP-Vangl2 from E15.5 wild-type control (*WT*) and K14-GFP-Vangl2 mouse epidermal lysates. Western blot using anti-GFP antibodies detects an ∼80KD band corresponding to the GFP-Vangl2 fusion protein (arrow). T=total protein input, IP=immunoprecipitate, Sup= supernatant. **(E)** Anti-GFP immunoprecipitates from wild-type control (*WT*) and K14-GFP-Vangl2 mouse epidermal lysates run on an SDS-PAGE gel stained with SPYRO-Ruby to detect total protein. The ∼80KD band present in the K14-GFP-Vangl2 but not wild type control immunoprecipitate was excised and processed for mass spectrometry. (**F)** Alignment of mouse Vangl2 with human, zebrafish, Xenopus, and *Drosophila* orthologs. The region spanning amino acids 269-312 of the C-terminal cytoplasmic tail is shown. Mass spectrometry analysis detected phosphorylation at highly conserved tyrosine residues 279 and 308 of mouse Vangl2 (bold with asterisk).

We furthermore detected Y-phosphorylation of Vang in a *fz*-null mutant background (Figure 1C), indicating that at least some Vang Y-phosphorylation is Fz independent, unlike the N-terminal Vang/Vangl2 S/T-cluster phosphorylation, which is Fz dependent and causes a detectable band shift on protein gels (see Figure 1C for loss of band shift; see also (Gao et al., 2011; Kelly et al., 2016; Strutt et al., 2019). Taken together, these *in vivo Drosophila* data suggest that Vang is phosphorylated on tyrosine residue(s) outside the defined Y341 residue and that at least some of these additional phosphorylation events might be Fz independent.

To gain insight into which tyrosines might be phosphorylated, we turned to a mass spectrometry-based approach. As it is technically challenging to obtain sufficient material from *Drosophila* pupal wings for such mass spectrometry studies (Yanfeng et al., 2011), we used mouse Vangl2 from embryonic skin. Due to its abundance and accessibility, the skin epidermis is an excellent model for biochemical analyses of proteins in their *in vivo* context. To identify post-translational modifications on epidermally expressed Vangl2, we performed an IP-MS analysis of GFP-Vangl2 protein purified from the skin of *K14-GFP-Vangl2* transgenic embryos (Devenport et al., 2011). Protein lysates were prepared from full thickness skin explants dissected from *K14-GFP-Vangl2* embryos at E15.5, and GFP-Vangl2 was immunoprecipitated using GFP-antibodies (Figure 1D). A prominent band of ∼80KD was present in immunoprecipitates from *K14-GFP-Vangl2* embryos but not wild type controls. This band was excised and processed for LC-MS/MS analysis (Figure 1E). Over 220 unique, high confidence peptides were recovered spanning 86% of the GFP-Vangl2 fusion protein. Importantly, phosphorylation modifications were detected at two different tyrosine residues Y279 and Y308 (Figure 1F), both of which are conserved across vertebrates and in *Drosophila* (see Figure 1F, and Suppl Fig S1D. Y279 is located within Vangl2’s TGN-sorting motif and is equivalent to Y341 in *Drosophila*, whose phosphorylation was previously implicated in Vangl2 trafficking (Guo et al., 2013; Kelly et al., 2016), indicating that our methods were able to detect functionally relevant Vang/Vangl2 phospho-tyrosine modifications (PTMs). Intriguingly, Y308 corresponds to residue Y374 in *Drosophila*, which is located within the previously defined Dsh/Dvl and Pk binding region of Vang (Jenny et al., 2003) (Figure 1F and 2A). The strong conservation of this tyrosine and its position within the Dvl and Pk binding region (see below) suggested that modification at this site may be important for Vang/Vangl function.

**Figure 2.**
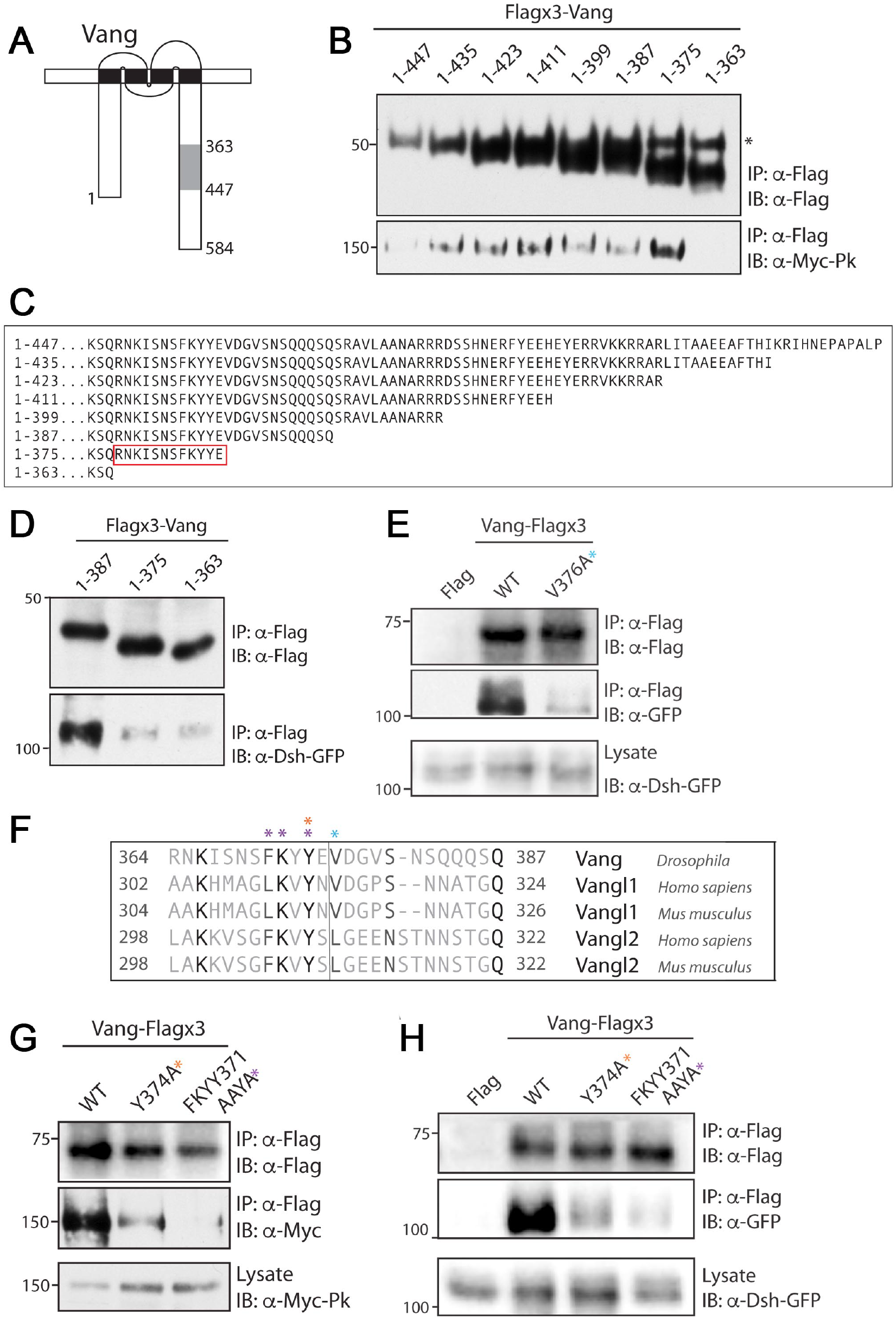
Pk and Dsh bind to adjacent, partially overlapping regions in the C-terminal tail of Vang. **(A)** Schematic of Vang showing the previously mapped region of Pk and Dsh binding along its C-terminal tail (shaded in gray), residues 363-447 (see Jenny et al. 2003). Y374 (the same residue as Y308 in mouse Vangl2) is indicated, note that it is located within the shaded region. **(B)** Western blot showing binding between Myc-Pk and C-terminal truncations of Flagx3-Vang, using a series of C-terminal truncations within the 363-447 stretch. Note binding is retained up until the 1-363 truncation, defining residues 363-375 in Vang as critical for its binding to Pk. (**C**) Schematic of the sequence of Vang C-terminal truncations as used in (B), red box highlights the amino acids required for Pk-binding. (**D**) Western blot showing binding between Dsh-GFP and C-terminal truncations of Flagx3-Vang. Note binding is retained up until the 1-375 truncation. (**E**) Western blot showing binding between Dsh-GFP and the V376A Vang-Flagx3 mutant. Note a marked reduction in binding was observed. V376 is at junction to the Pk-binding region (and largely conserved), suggesting an overlap in binding regions between Pk and Dsh. **(F)** Sequence alignment showing the conservation of amino acids in the *Drosophila* Vang 364-387 region. Colored asterisks highlight the amino acid mutated in binding experiments shown in panels E, G and H. **(G)** Western blot showing binding between Myc-Pk and selected Vang-Flagx3 mutants as indicated. Colored asterisks highlight specific amino acids in sequence schematic in (**F**) mutated in the experiment. Note marked reduction in binding for the single mutant (Y374A) and almost complete loss of binding in the triple mutant FKYY371AAYA. **(H)** Western blot showing binding between Dsh-GFP and the indicated Vang-Flagx3 mutants. Note the reduction in binding follows a similar pattern to what was observed with Pk (compare to panel **G**). Importantly, the equivalent Y residue to Y374 is detected as phosphorylated in mouse Vangl2 (see Fig. 1D-G).

### Vang binds Pk and Dsh at an overlapping region around the Y374/Y308 phospho-site

Vang/Vangl proteins have been shown to physically interact *in vitro* with all other members of the core PCP factor group as mentioned above. The molecular interactions of Vang with the cytoplasmic core factors have previously been mapped to specific regions within the C-terminal tail of the protein (Jenny et al., 2003). In particular, previous work in *Drosophila* has demonstrated that Vang amino acids 363-447 are required for its interaction with Pk and Dsh (Jenny et al., 2003). Strikingly, as mentioned above and defined in the MS studies, one of the conserved PTMs coincided with this region (shaded C-tail stretch in Figure 2A). We thus aimed to refine the binding sites of Pk and Dsh within this region and define the potential role of the phosphorylated Y-residue in this context. We then wished to potentially generate specific binding mutants in Vang/Vangl to modulate its interactions with cytoplasmic core factors, with the ultimate goal to assess their functional importance *in vivo*.

A key binding partner of Vang/Vangl proteins is Pk, with the two proteins co-localizing together stably during the mature stages of PCP establishment, with these asymmetrically localized complexes forming as a result of interactions and transition stages/complexes among the core PCP genes (Jenny et al., 2003). To refine the binding site of Pk we made a series of C-terminal truncations of *Drosophila* Vang covering the known binding region, amino acids 363-447 in the C-terminal tail, and reduced this binding domain by 12 amino acids with each truncation (Figure 2B-C). Pull-down experiments, performed in S2 cells, revealed that Pk could interact with all C-terminal Vang truncations with the exception of 1-363, which suggested that amino acids 364-375 were required for Pk-binding (Figure 2B, see 2C for sequence schematic of deletion constructs). Importantly, this amino acid stretch contains the conserved PTM site, and an FKxY motif that was identified as phosphorylated *in vivo* in the mass spec studies (see above).

We next assessed Vang interactions with the other cytoplasmic effectors that were also previously shown to interact within the C-tail region (Jenny et al., 2003; Jenny et al., 2005). To refine these binding sites, we again turned to the C-terminal truncations set, now to assess Dsh interaction(s). Strikingly, we observed that Dsh also showed markedly reduced binding when residues 363-387 were deleted (Figure 2C-D). In the case of Dsh we observed binding loss with the 1-375 truncation, directly adjacent to the Pk binding area, suggesting that amino acids 376-387 are critical for the Vang-Dsh interaction (Figures 2C-D). As this region was immediately adjacent to the Pk binding site, we postulated that the binding site for Dsh may be extended and could require additional amino acids. Analysis of the conservation of the amino acids within 376-387 revealed a partially conserved valine (V) at position 376 (Figure 2E-F). Indeed, a V376A substitution led to a marked reduction in Dsh binding (Figure 2E-F). In contrast, we did not observe an impact of the V376A substitution on the Vang-Pk interaction (Suppl Fig. S2). Taken together, these data suggest that Pk and Dsh bind to regions immediately adjacent to each other, and likely overlapping.

As, again, this region of Vang/Vangl2 contains a conserved PTM at Y374 in *Drosophila* or Y308 in mouse Vangl2 (see Figure 2F for sequence comparison), we tested whether this Y-residue and/or associated motif affects interaction with either Pk or Dsh or both. Strikingly, a Vang-Y374A substitution caused a markedly diminished binding of both Pk and Dsh (Figure 2G and H, respectively). Importantly, this effect was further enhanced in the triple mutant affecting the whole FKxY motif (FKYY371AAYA) with interaction of either Pk or Dsh further reduced (Figure 2G and H, most right lanes each, respectively). It is worth noting that in control experiments with the core PCP effector Dgo or the A/B polarity protein Scribble (Scrib), with both demonstrated to interact with the C-tail of Vang/Vangl2 (Courbard et al., 2009; Das et al., 2004; Jenny et al., 2005; Montcouquiol et al., 2003), these mutations in the FKYY motif of Vang did not display any effect on their binding efficiency to Vang (Suppl. Figure S2). Furthermore, the specificity and importance of Y374 and the FKYY motif for interaction with Pk and Dsh within the defined binding region was confirmed via additional point mutations in the same region, as for example mutations of K366, K372, or F372 did not demonstrate a detectable requirement for Pk binding (Suppl. Figure S2A). Similarly, V376 was specifically required for the Dsh interaction and did not affect the Vang-Pk binding (Suppl. Figure S2B).

Taken together, these data suggested that Pk and Dsh, two antagonistic cytoplasmic core PCP factors, share an overlapping binding site centered around a conserved PTM (Y374 in *Drosophila* and Y308 in mouse Vangl2) in the C-terminal tail region of Vang proteins with both Pk and Dsh requiring Y374 for binding within a conserved motif. In addition, our data also suggested that we have defined a specific point mutant at V376 that modulated Dsh-binding but did not have an effect on Pk (see also below).

### The phosphorylation/charged state at Y374 contributes to binding regulation

Previous work in *Drosophila* and mouse has demonstrated that serine/threonine phosphorylation of Vang/Vangl2 regulates its function (Gao et al., 2011; Kelly et al., 2016; Strutt et al., 2019). As the key residue responsible for binding to both Pk and Dsh was a phosphorylated tyrosine, we thus looked to test whether phosphorylation, or the charge associated with phosphorylation, at this site might regulate binding.

As such, we performed substitution of Y374 to phenylalanine (F) to assess binding to each effector. The substitution Y374F retains the structural aromatic ring associated with tyrosine, but as it cannot be phosphorylated it remains an uncharged residue. Importantly, this also allowed comparison to the Y374A substitution, shown to diminish binding of both Pk and Dsh (see above, Figure 2G-H). Strikingly, we observed that Pk showed equally diminished binding to both Y374A and Y374F (Figure 3A), while Dsh retained binding to Y374F (Figure 3B; Dsh-binding was reduced in the Y374A substitution in the same experiment as a control, see above). These data suggested that Dsh binding has a preference for the structure of the amino acid, an aromatic ring, shared between tyrosine and phenylalanine, but does not seem to care whether the residue can be phosphorylated (charged). To ask more directly whether a charge at Y374 can contribute to binding regulation, we tested a Y374D substitution (with D, a charged residue, to possibly mimic a phosphorylation charge). Interestingly, Pk displayed significant binding to Y374D, while Dsh did not (Figure 3C and D, respectively; compare to wt-control and Y374F in same panels). To increase the charge (and possibly be closer to a phospho-mimetic situation), we also generated YY373DD substitutions with two charged residues in the positions 373-374. Strikingly, the “phosphomimetic” Vang YY373DD bound to Pk better than the single Y374D substitution, suggesting that indeed a strong charge at Y374 (via phosphorylation) is the preferred state for Pk binding (Figure 3C). In contrast, there was no significant effect on the Dsh interaction as compared to Y374D (Figure 3D). These data suggested that Dsh requires an aromatic ring at the binding region centered on Y374 and does not require phosphorylation, while Pk-binding appeared to be dependent on charge and thus phosphorylation.

**Figure 3.**
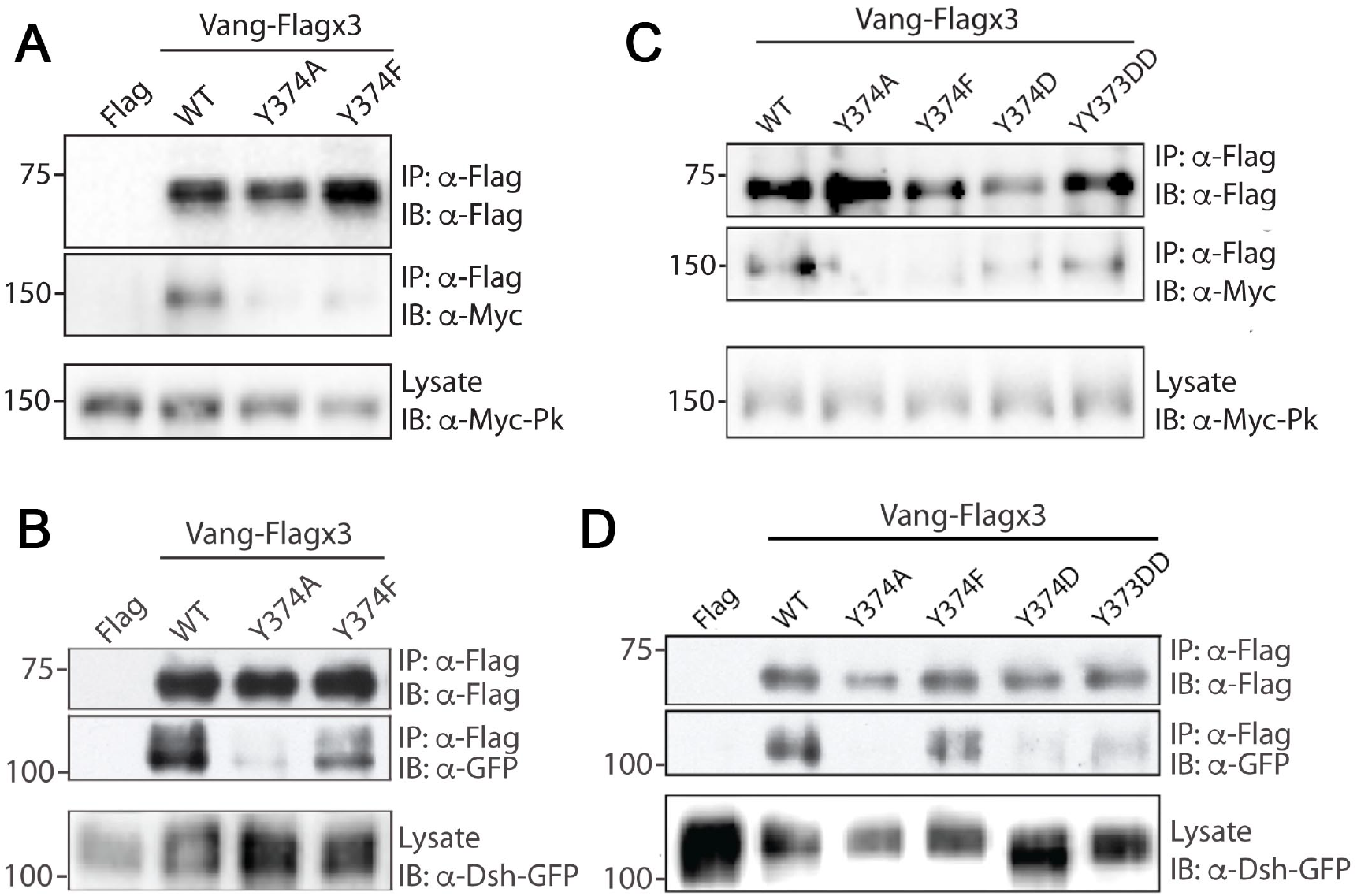
Differential substitutions at Y374 reveal a role for charge/phosphorylation in binding regulation. (**A**) Western blot showing binding between Myc-Pk and the indicated Vang-Flagx3 mutants, Y374A and Y374F. Note binding is markedly reduced in both cases. (**B**) Western blot showing binding between Dsh-GFP and the indicated Vang-Flagx3 mutants. Note binding is reduced with an alanine substitution, Y374A, but not with phenylalanine, Y374F, which retains the aromatic ring feature of tyrosine. (**C**) Charged amino acid substitutions of Y374 partially restore Pk binding. Western blot showing binding between Myc-Pk and the indicated Vang-Flagx3 Y374 substitutions. Note that Y374D and YY373DD recover significant binding of Pk, as compared to Y374A and Y374F. (**D**) Charged amino acids do not enhance Dsh interaction with Vang. Western blot showing binding between Dsh-GFP and the indicated Vang-Flagx3 substitutions. Note that Vang Y374F is by far the best interacting mutant, again confirming a requirement of an aromatic ring feature/structure for Dsh binding (see also panel **B**).

To confirm and corroborate this hypothesis, we also generated the equivalent binding region as an *in vitro* peptide and phospho-peptide. The peptide generated encompasses the Vang region 367-381 that should be sufficient to bind to both Pk and Dsh, in its phosphorylated form at Y374, ISNSFKY[pY]EVDGVSN-amide (pY-peptide, Suppl. Fig. S3A) and as a non-phosphorylated control (Y-peptide, Suppl. Fig. S3A). Importantly these peptide interaction assays could confirm that the binding region defined above (Figure 2) is indeed sufficient for effector binding, not just necessary, and that actual phosphorylation matters for the Pk interaction. Consistent with the co-IP studies described above, the (phosphorylated) pY-peptide preferentially interacted with Pk (Suppl. Figure S3B), whereas the un-phosphorylated control peptide (“Y”) was preferentially bound by Dsh (Suppl. Figure S3C). To further corroborate the binding preferences, we assayed interactions with purified proteins in a competitive binding assay (Suppl. Fig. S3D). Strikingly, Dsh-GFP protein when bound to the pY peptide (coupled to beads) was readily outcompeted by GFP-Pk (Suppl. Figure S3D, quantified in Fig. S3E), whereas Dsh bound to the control “Y”-peptide was not competed away by Pk (Suppl. Figure S3D).

In summary, all the above data taken together are consistent with the model that (i) the Vang region centered around Y374 is sufficient for binding to Pk and Dsh, (ii) Pk preferentially interacts with this binding region when Y374 is phosphorylated, and (iii) Dsh requires the structural features of an aromatic ring in Y374, and not phosphorylation in its binding behavior to Vang/Vangl. Furthermore, these experiments defined a single amino acid mutation, Y374F, that selectively abrogated Vang-Pk binding, but retained the Vang-Dsh interaction (see above), and VangV376A shows the opposite, maintaining Pk interaction but causing a loss of the Vang-Dsh binding. These observations also allowed for the first time to functionally test for a Vang-Dsh binding requirement *in vivo* (see below).

### Single point mutants show PCP defects

To investigate the functional consequence *in vivo* of tempering the interaction of Vang with Pk and Dsh, we performed rescue experiments using transgenes carrying our different single point mutants. For this, we expressed Vang-Flagx3 WT, Y374A, Y374F and V376A, using a direct *tubulin*-driven expression in a *Vang* null mutant background (*Vang*^*6/6*^). The control construct *tub-VangWT* was able to fully rescue cellular orientation in *Drosophila* wings (Figure 4A-C). In contrast, each of the single point mutants, Vang-Y374A, Vang-Y374F and Vang-V376A, failed to rescue the *Vang*^*6*^ loss-of-function phenotype, displaying varying degrees of cellular hair orientation defects (Figure 4C, quantified in 4D). Consistent with the molecular binding data, Vang-Y374A, which does not bind either Pk or Dsh showed the strongest defects largely resembling the *Vang* null phenotype. The point mutations that lead to loss of binding to a single effector (Y374F only binding Dsh, and V376A only binding to Pk) showed a weaker phenotype as compared to Y374A (Figure 4C-D). This was further supported through quantification of cellular hair orientation, using the FijiWingsPolarity plug-in (Dobens et al., 2018), that revealed a significant variation from the *Vang WT* control in each case, and also between Vang-Y374A and the other point mutations (Figure 4C-D). Importantly, the difference in phenotype was not due to variation in expression levels, as all genotypes showed equivalent protein levels in larval disc lysates (Figure 4E).

**Figure 4.**
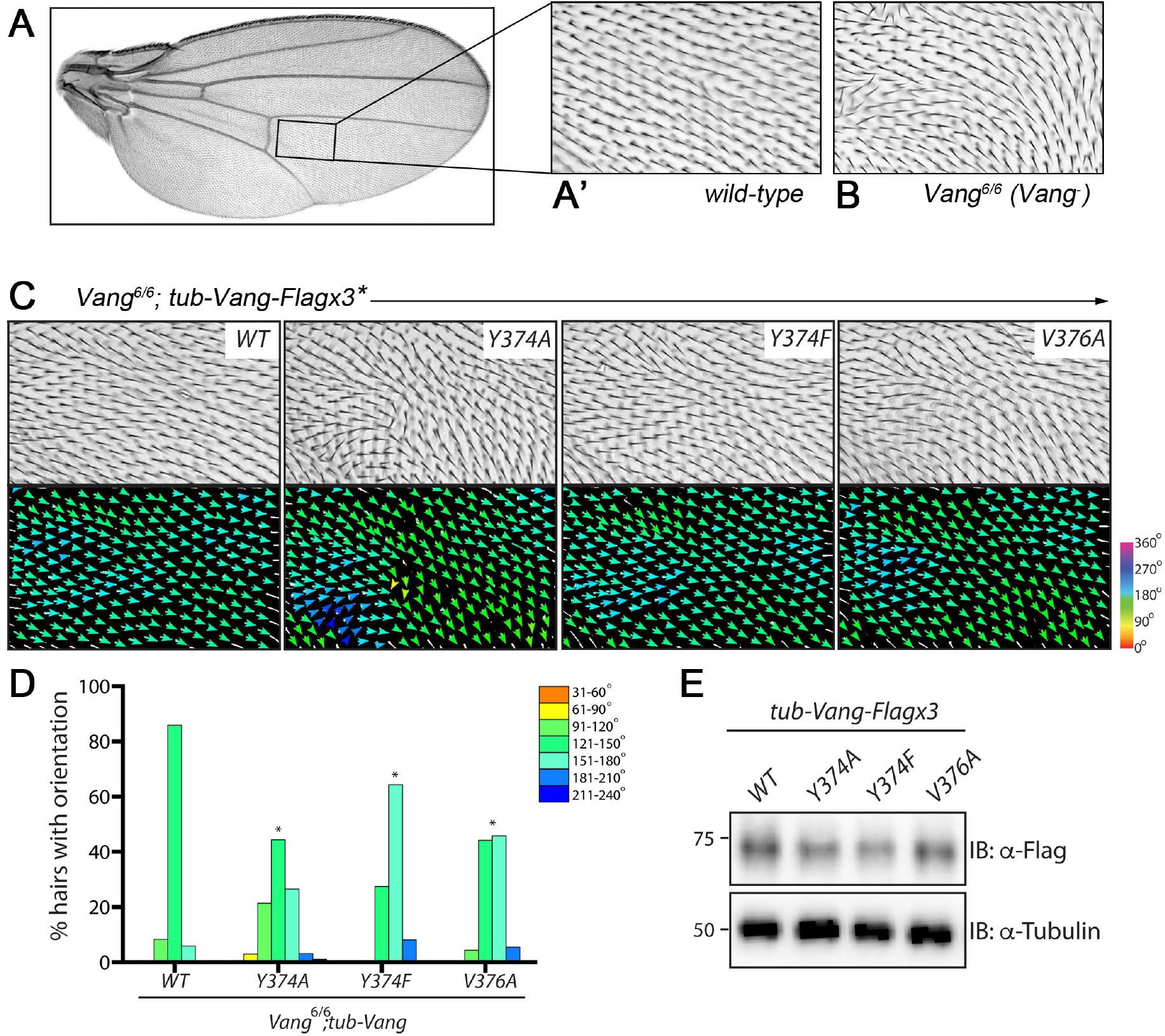
Single Vang point mutants in the Pk and Dsh interaction region display PCP defects *in vivo*. (**A**) *Drosophila* wing (distal is right) and (**A’**) higher magnification of boxed area in wing on left. Note cellular orientation being reflected by single actin hairs pointing all distally. (**B**) Sample image of same region as in **A’** of *Vang*^*6/6*^ (null mutant) wing. Note misoriented cellular hairs forming waves and whorls in the *Vang*^*-/-*^ background. (**C**) Sample images of cellular (re)orientation in adult wings of *Vang* mutant flies (same region as in **A’-B**) upon rescue of *Vang*^-/-^ animals with *tubulin-*promoter expressed *Vang-Flagx3 WT (wild-type), Y374A, Y374F* or *V376A*, respectively as indicated. Angles of cellular orientation are visualized through colored arrows in lower panels (see scale on right). Note that while Vang-*WT* fully rescues the mutant phenotype (and appears like wild-type, compare to **A’**), all single point mutants fail to rescue the mutant defects. Note also that Vang-Y374A resembles the *Vang* null phenotype (cf. to panel **B**), the other point mutations, display weaker phenotypes. (**D**) Quantification of the percentage of cellular actin hairs oriented within a specific angle range for the indicated genotypes. Angles were determined using plugin FijiWingsPolarity (Dobens et al. 2018). Cellular hair orientation angles from 3 independent wings were combined, and data binned for statistical analysis via Chi-squared test. *p*<0.001 is indicated by * as compared to the rescue of Vang-*WT*. (**E**) Western blot of wing disc lysates from the indicated genotypes. Note similar levels of expression in each transgene, mutant.

To confirm the different *in vivo* behavior of the individual Vang binding mutants, we also tested their effects in a gain-of-function (GOF) assay in developing wings. A classic feature of core PCP factors is that their overexpression, GOF scenario, also causes PCP defects. Each core factor has not only a reproducible LOF phenotype, but also a GOF phenotype that causes stereo-typic defects highly similar from wing to wing of any given genotype. As such the Vang-*wt* GOF (via *nubbin-Gal4*) induces similar misorientations from wing to wing (in any specific region of a wing; see ROI samples in Suppl. Fig. S4A-B, boxed areas in blue and red in S4A, correspond to left and right panels in B, respectively). In each case when Vang is expressed via *nubbin-Gal4*, Vang-*wt* or any Vang-mutant (see indicated mutant genotypes in Fig. S4B-C), cellular polarity is perturbed and while wings of one genotype look similar, they have a different appearance between distinct Vang-mutant genotypes (Supp. Fig. S4B). This allowed us to ask whether, for example, a Vang-Y374F (binding only Dsh) is distinct from a Vang-YY373DD (binding predominantly Pk) or Vang-Y374A (binding neither factor). Strikingly, all specific Vang mutations tested displayed distinct behavior (Fig. S4B, see quantifications in angle distribution graphs on right of each panel), although the levels of each mutant protein were comparable (Suppl. Fig. S4C). The notable exception was when Vang-YY373DD and Vang-V376A were compared (Suppl. Fig S4B bottom two rows, which appeared to cause similar reorientations. However, this effect was consistent with all data above, as both these mutant Vang isoforms bind Pk only.

Combined together with the rescue data (Figure 4), all these results are consistent with the notion that Vang is required to interact with/bind to both cytoplasmic PCP core factors, Pk and Dsh, during the establishment of planar polarity *in vivo*. Together with the biochemical dissection, these data thus suggest that Y374 phosphorylation is critical for the physiological activity of Vang in the process of PCP establishment and the resolution of the core PCP genes into the Vang and Fz associated antagonistic complexes (see also Discussion).

## Discussion

Here we provide *in vivo* evidence that tyrosine phosphorylation on a conserved tyrosine (Y374 in *Drosophila* Vang and Y308 in mouse Vangl2) within the cytoplasmic C-terminal tail of Vang/Vangl is required for its function and regulates its interaction(s) with the core PCP factors Pk and Dsh. Through mass spectrometry analyses with mouse skin epidermis samples we identified phosphorylation on Y308 in mouse Vangl2 (equivalent to Y374 in *Drosophila* Vang). Strikingly, this tyrosine coincides with the overlapping binding regions of Pk and Dsh in Vang/Vangl, and, importantly, its phosphorylation status regulates selective binding between Pk and Dsh, with phosphorylation tipping the balance towards Pk binding. We also demonstrate *in vivo* that binding of Vang to both cytoplasmic core PCP factors is physiologically important (which is the first *in vivo* evidence for a Vang-Dsh binding requirement). Our study provides novel insight into the critical importance of Vang tyrosine phosphorylation and reveals mechanistic features of how regulation of the binding of antagonistic PCP factors to Vang/Vangl during the process of PCP complex segregation and polarity establishment is achieved.

### Phosphorylation of Y374/Y308 at the center of Vang/Vangl interactions with Pk and Dsh

While previous work defined a broad region within the C-tail of Vang to interact with both Pk and Dsh (Jenny et al., 2003; Jenny et al., 2005), the mechanistic regulation and physiological significance of these interactions remained unresolved. Importantly, the defined region is conserved between *Drosophila Vang* and mammalian *Vangl1/2* genes. Our data reveal that a small stretch of amino acids within this broader region is both necessary (as shown in the whole Vang protein) and sufficient (as deduced from the *in vitro* peptide assays) to interact with both cytoplasmic core PCP factors. This peptide/region is well conserved between all Vang family members and centered on the tyrosine, which can be phosphorylated, as our mass spec data reveal. Mutational studies define that Pk binding is mediated by tyrosine phosphorylation and associated charge, while Dsh requires the aromatic ring found in tyrosine (and also phenylalanine) for its binding to Vang. It is worth noting that this binding region, shared by both Pk and Dsh for binding to Vang, is specific for these two factors, as other cytoplasmic Vang associated PCP proteins, for example, Dgo and Scrib, are not affected by any mutations within this domain.

While the importance of the Vang-Pk complex has been well documented *in vivo* and is also the core of one of the two stable PCP core complexes that result from PCP factor interactions and signaling, defining cellular polarity (reviewed in (Adler, 2012; Axelrod, 2020; Butler and Wallingford, 2017; Devenport, 2014; Goodrich and Strutt, 2011; Humphries and Mlodzik, 2018; Yang and Mlodzik, 2015), an interaction between Vang and Dsh/Dvl family members has only been shown biochemically (Bastock et al., 2003; Jenny et al., 2003; Park and Moon, 2002; Seo et al., 2017; Torban et al., 2004). The detailed dissection of binding requirements allowed us to generate single point mutations in Vang that separate Vang binding to only one of these cytoplasmic factors, either Pk or Dsh. The associated *in vivo* rescue constructs provided the possibility for physiological testing of a functional requirement of the individual interactions between Vang and Dsh or Pk. A point mutation interfering with binding of both factors to Vang showed very little detectable rescue activity (with strong similarity to the *Vang*^*-*^ null phenotype), suggesting that much of the Vang function is mediated by interaction with Pk and Dsh. Strikingly, point mutations affecting individual Vang-Pk or Vang-Dsh interactions also failed to rescue the *Vang*^-/-^ defects to wild-type. In both cases, only a partial rescue was observed, indicating that interactions with either cytoplasmic PCP factor, Pk and Dsh, are critical for Vang function in PCP establishment. As such this is the first physiological *in vivo* evidence demonstrating that Vang has a requirement to interact with Dsh.

### How does Vang Y374/Y308 phosphorylation affect PCP complex resolution?

It is intriguing to think about how phosphorylation, and lack thereof, affects the formation of stable core PCP complexes. Our data indicate that binding of Dsh/Dvl to Vang/Vangl is physiological and important, and yet in standard co-localization studies Vang-Dsh complexes are not detected. How does Dsh binding to an unphosphorylated Y374 region affect core PCP complex formation? There are a few plausible scenarios. First, a Vang-Pk interaction, which we assume is stable in the correct region and context of a cell, should likely not form in areas of the cell membrane where it is not supposed to end up. As such a Vang-Dsh transient/intermediary interaction might serve a function to prevent such (stable) binding. If the kinase is localized to, or active, in the correct area, then - and only then - a switch from Vang-Dsh to a stable Vang-Pk interaction could occur. As such, Dsh binding to this Vang region might also prevent the kinase to act on Vang prematurely and hence prevent formation of a stable Vang-Pk complex in the wrong region of a given cell. In such a mechanistic scenario Dsh would keep Vang/Vangl “flexible” to find the right cellular context, where and when the presence of the right kinase would cause the switch to a Vang-P at Y374 (Y308 in mVangl2) and force a stable interaction with Pk and its local effectors. While this is an intriguing mechanistic proposition, it remains a speculation and complex *in vivo* experiments might be necessary to proof this.

### What is the tyrosine kinase acting on Vang/Vangl?

It is currently unclear which tyrosine kinase(s) act on Vang to mediate its phosphorylation on Y374 (Y308 in mVangl2). Sequence motif searches with the site suggest that Src family kinases could be involved, and no other kinase has a higher probability by motif search to act in this context. It is however technically very difficult to prove that Src kinases indeed act on Vang and this site *in vivo* (see below), and *in vitro* assays are largely meaningless as any tyrosine kinases tested could phosphorylate Vang in such assays. In attempted studies *in vivo*, redundancy of Src kinases is an issue in both our systems, mouse skin and *Drosophila* wings (there are several Src family kinases in both *Drosophila* and mice, for example (Lowell et al., 1994; Pedraza et al., 2004; Simon et al., 1983; Stein et al., 1994; Thomas et al., 1995)), in addition to the cell survival requirements, many cellular functions are associated with Src family kinases (rev in e.g. (Chen et al., 2018; Espada and Martin-Perez, 2017; Sirvent et al., 2020)). For example, in *Drosophila* the two main Src family members are either viable with no overt developmental phenotype in imaginal discs (Src64, redundancy) or are cell lethal (Src42) when analyzed *in vivo* (Pedraza et al., 2004; Simon et al., 1983). They have also been linked to a vast variety of cellular functions, ranging from cytoskeletal regulation, cell adhesion, synaptic plasticity, proliferation, cell death, and others (rev in. (Chen et al., 2018; Espada and Martin-Perez, 2017; Sirvent et al., 2020)). Src kinases remain nonetheless likely candidate(s), as we (i) observe GOF phenotypes consistent with a PCP function and (ii) genetic interactions with these Src GOF defects suggest that Vang is required in these contexts. However, again, a loss-of-function scenario to really show a Src function in PCP establishment remains elusive and will be the focus of future studies.

## Acknowledgements

We are grateful to Andreas Jenny and Ursula Weber for plasmids, and we thank all Devenport and Mlodzik lab members, past and present, for helpful input and discussions. We thank Saw Kyin and David Perlman of the Molecular Biology Proteomics & Mass Spectrometry Core facility at Princeton University who provided technical support and analyzed the mass spectrometry data. This research was supported by National Institutes of Health grants R01 AR066070 (to D.D.) and R35 GM127103 (to M.M.), and also in part by an EMBO post-doctoral fellowship to A.C.H., as well as an New Jersey Commission on Cancer Research post-doctoral fellowship to C.C.H.

## Materials and Methods

### DNA constructs and S2 culture

Constructs used were as follows; pAc5.1-Flagx3, pAc5.1-Myc-Pk, pAc5.1-Dsh-GFP, pAc5.1-Scrib-PDZ-3-4-HA and pAc5.1-HA-Dgo (all gifts from Dr. Jenny, AECOM, USA) and pAc5.1-Vang-Flagx3 (Humphries et al., 2020). GFP-TZ was used as a control, and is the *Drosophila* ciliary protein Mks1. pAc5.1-Vang-Flagx3-Y374A, FKYY371AAYA, Y374F and V376A were generated though site-directed mutagenesis. C-terminal truncations of Vang were generated through PCR and cloned into pAc5.1-Flagx3 using NotI-XbaI restriction sites. All primers used are available upon request.

Unless otherwise stated, lysates were prepared from S2 cells. S2 cells were maintained according to standard protocols, and were grown in Schneider’s Medium (Gibco) supplemented with 10% heat-inactivated Fetal Bovine Serum (Gibco). Cells were plated in 12 well plates at a dilution of 1.5×10^6^ and were transfected with the indicated constructs using Effectene (Qiagen) according to manufacturer’s protocols. Cells were lysed ∼48 hrs later in buffer containing 50mM Tris-HCl pH 7.5, 150mM NaCl, 1mM EDTA and 1% Triton-X-100.

### Pull-downs and immunoblotting

For Flag pull-down experiments, lysates were extracted from S2 cells or from larval wing discs and were incubated at 4**°**C overnight with 10μl anti-Flag M2 affinity gel per sample. Washes were performed in buffer containing 50mM Tris-HCl pH 7.5, 350mM NaCl, 1mM EDTA, 0.1% SDS before elution in final sample buffer. For peptide binding, Vang peptides were conjugated to beads to enable assessment of direct binding. Lysates were generated in S2 cells, and the lysate was divided equally between tubes containing either 20μl of conjugated phospho-peptide or the unphosphorylated form and incubated at 4**°**C overnight. Washes were performed in buffer containing 10mM Tris-HCl pH7.5, 350mM NaCl, and 0.5mM EDTA before elution in final sample buffer. For GFP pull-downs. Samples were resolved by polyacrylamide gel electrophoresis according to standard protocols. The following primary antibodies were used for immunoblotting; Flag (Sigma M2 1:5000), Gamma-tubulin (Sigma GTU-88 1:1000), GFP (Roche 7.1&13.1 1:1000), Myc (SCBT 9E10 1:1000), Phospho-tyrosine (Millipore PY20 1:1000).

### Drosophila strains, dissections and phenotypic analyses

Flies were raised on standard medium, and maintained at 25**°**C unless otherwise stated. To generate *UAS-Vang-Flagx3* mutant transgenic flies, site-directed mutagenesis was performed on vector pUAST-Vang-Flagx3-attB (Humphries et al. 2020), before insertion into BDSC stock number 9750. To generate *tub-Vang-Flagx3* mutant flies, site-directed mutagenesis was performed on vector pCaSpeR-tub-Vang-Flagx3 (Kelly et al. 2016). All transgenic strains were generated by Bestgene Inc.

Wing discs were dissected from third instar larvae and prepared through incubation in final sample buffer at 95**°**C. For phosphatase treatment, wing discs were collected in PBS and transferred to lysis buffer containing 50mM Tris-HCl pH 7.5, 150mM NaCl, 1mM EDTA, and 1% Triton-X. Lysates were then incubated at 30**°**C for 30 minutes with lambda protein phosphatase (NEB).

Adult wings were collected in PBS containing 0.1% Triton-X-100 (PBST) and incubated for 1 hr at room temperature before mounting in 80% glycerol in PBS. Hair orientation was quantified using the FijiWingsPolarity plugin (Dobens et al., 2018). The quantification was performed on equivalent regions of 3 wings per genotype, and angles binned into 30° degree segments to allow for statistical analysis via a Chi-squared test.

### Mouse lines and embryonic skin harvesting

K14-GFP-Vangl2 transgenic mice (FVB background, (Devenport et al., 2011)) were housed in an AAALAC-accredited facility following the Guide for the Care and Use of Laboratory Animals. Animal maintenance and husbandry followed the laboratory Animal Welfare Act. Princeton University’s Institutional Animal Care and Use Committee (IACUC) approves all animal procedures. E15.5 embryos were harvested from K14-Vangl2-GFP heterozygous dams in cold PBS and screened for GFP expression using a stereomicroscope equipped with epifluorescence. Full thickness backskins were dissected from both GFP+ and GFP-littermates embryos and flash frozen immediately in liquid nitrogen (LN2). LN2 was removed by evaporation and frozen backskins were stored for up to 3 months at -80C until cryolysis.

### Epidermal cryolysis and immunoprecipitation of GFP-Vangl2

Frozen skin samples pooled from four (for IP-Western) or eight (for IP-MS) GFP-Vangl2 and control backskins were processed via cryolysis using a CryoMill (brand type etc). Briefly, 2ml LN2-frozen lysis buffer droplets (Tissue Extraction Buffer, 1% Triton X-100, 10mM EDTA, 0.3mg/ml PMSF plus protease inhibitors in PBS) were mixed with frozen skin samples and processed by cryogenic grinding for 20 min using a ball mill cooled with LN2. Finely ground frozen epidermal-lysis buffer mixtures were lysed by thawing on ice for 1.5-2hrs. Lysates were cleared by adding 50ul Pansorbin and centrifuging at 14K rpm for 10min.

Pre-cleared lysates were transferred to pre-washed, αGFP (rabbit αGFP, AbCam) antibody bound beads and incubated for 3 hours at 4 degrees C. Immunoprecipitates were washed twice with lysis buffer and eluted with 40ul 2X SDS sample buffer. For western blotting, 40ul total lysate and 40ul IP were run on a 10% SDS-PAGE gel and transferred to a PDVF membrane, blocked for 1hr at room temperature, incubated with primary, chicken αGFP (1:5000, Abcam) overnight at 4°C. After several washes in PBS-T, membrane was incubated for 45 min at room temperature with HRP α-chicken secondaries (1:2500), developed with BioRad Clarity ECL reagent and imaged on both film and with BioRad imager.

For IP-MS samples, 20ul total lysate and 40ul IP were run on a 7.5% SDS-PAGE gel. Gel was fixed in 50% MeOH + 7% Acetic Acid for 30-60 minutes, rinsed with H_2_0, stained with SPYRO Ruby overnight, and washed twice for 5 min each in 10% MeOH + 7% Acetic Acid. Gel imaging was performed on a Typhoon FLA-7000 (GE Healthcare).

### Proteomic sample preparation and mass spectrometry

IP Bands at ∼85-90KD were excised then diced, and subjected to in-gel thiol reduction/alkylation and trypsin digestion using a method adapted from Shevchenko et al (2006) to process samples for LC-MS/MS. Briefly, gel cubes were destained and washed extensively in 100 mM ammonium bicarbonate buffer, pH 8.8 (ABC), treated with 50 mM TCEP in ABC for 1 h at 55°C, washed, subjected to alkylation with 55 mM iodoacetamide in ABC for 30 min at room temperature in the dark, washed, and finally digested overnight with 1 ug Promega Trypsin Gold (Promega) per gel slice. Peptides from in-gel digest eluates were desalted using STAGE-Tips 9 prior to LC-MS analyses.

LC-MS/MS analyses were performed on a high-resolution, high-mass-accuracy, reversed-phase nano-UPLC-MS platform, consisting of an Easy nLC Ultra 1000 nano-UPLC system coupled to an Orbi Elite mass spectrometer (ThermoFisher Scientific) equipped with a Flex Ion source (Proxeon Biosystems, Odense, Denmark). LC was conducted using a trapping capillary column (150 µm x ca. 40 mm, packed with 3 µm, 100 Å Magic AQ C18 resin, Michrom, Auburn, CA) at a flow rate of 5 µL/min for 4 min, followed by an analytical capillary column (75 µm x ca. 45 cm, packed with 3 µm, 100 Å Magic AQ C18 resin, Michrom) under a linear gradient of A and B solutions (solution A: 3% acetonitrile/ 0.1% formic acid; solution B: 97% acetonitrile/ 0.1% formic acid) from 5%-35% B over 90 at a flow rate of 300 nL/min. Nanospray was achieved using Picospray tips (New Objective, Woburn, MA) at a voltage of 2.4 kV, with the Elite heated capillary at 275°C. Full-scan (m/z 335–1800) positive-ion mass spectra were acquired in the Orbitrap at a resolution setting of 120,000. MS/MS spectra were simultaneously acquired using CID in the LTQ for the fifteen most abundant multiply charged species in the full-scan spectra, having signal intensities of >1000 NL. To aid in phosphosite mapping, Vangl2-positive slices were subjected LC-MS/MS over a 180 min gradient using the CID parameters above and also a second round of LC-MS/MS over a 180 min gradient during which MS/MS spectra were acquired by multistage activation (MSA) for the top 10 most abundant ions in the full-scan spectra, using excitation at the precursor *m/z* value as well as those corresponding to the neutral losses of phosphonic and phosphoric acids for ions of charge +2 and +3. Lockmass was employed, maintaining calibration to 2-3 ppm of accurate mass.

### Mass Spectrometric Data Analysis

Resultant LC-MS/MS raw data files were processed using ProteomeDiscoverer (v. 1.4, ThermoFisher), to match MS/MS spectra against the UniProt *Mus musculus* database, or a GFP-Vangl2 fusion protein construct subset database using the Mascot search engine (v. 2.4, Matrix Science, London, UK.), allowing for a parent ion mass window of ±6 ppm, ≤ 3 missed trypsin cleavages, serine, threonine and tyrosine phosphorylation, methionine oxidation, asparagine and glutamine deamination and *N*-terminal protein acetylation as variable modifications, and carbamidomethylation of cysteines as a fixed modification. Peptide assignment cut-offs were specified at a high confidence level (<1% FDR). Phosphosite localization confidence scoring was achieved using the PhosphoRS 9 (Taus et al., 2011) (v. 3.1) node within the ProteomeDiscoverer framework. Relative abundance levels for proteins between experimental and controls were estimated using spectral counting. Raw mass spectra were visualized using Xcalibur (v. 2.2, ThermoFisher) and peptide spectral matches were visualized using ProteomeDiscoverer or Scaffold (v. 4.3.4, Proteome Software, Portland, OR). All phosphopeptide assignments were further validated by manual inspection.

## References

Adler, P.N. (2012). The frizzled/stan pathway and planar cell polarity in the Drosophila wing. Curr Top Dev Biol 101, 1–31.

Axelrod, J.D. (2020). Planar cell polarity signaling in the development of left-right asymmetry. Curr Opin Cell Biol 62, 61–69.

Bastock, R., Strutt, H., and Strutt, D. (2003). Strabismus is asymmetrically localised and binds to Prickle and Dishevelled during Drosophila planar polarity patterning. Development 130, 3007–3014.

Boutin, C., Labedan, P., Dimidschstein, J., Richard, F., Cremer, H., Andre, P., Yang, Y., Montcouquiol, M., Goffinet, A.M., and Tissir, F. (2014). A dual role for planar cell polarity genes in ciliated cells. Proc Natl Acad Sci U S A 111, E3129–3138.

Butler, M.T., and Wallingford, J.B. (2017). Planar cell polarity in development and disease. Nat Rev Mol Cell Biol 18, 375–388.

Cai, C., and Shi, O. (2014). Genetic evidence in planar cell polarity signaling pathway in human neural tube defects. Front Med 8, 68–78.

Carvajal-Gonzalez, J.M., Mulero-Navarro, S., and Mlodzik, M. (2016). Centriole positioning in epithelial cells and its intimate relationship with planar cell polarity. Bioessays 38, 1234–1245.

Cetera, M., Leybova, L., Joyce, B., and Devenport, D. (2018). Counter-rotational cell flows drive morphological and cell fate asymmetries in mammalian hair follicles. Nat Cell Biol 20, 541–552.

Chen, W.S., Antic, D., Matis, M., Logan, C.Y., Povelones, M., Anderson, G.A., Nusse, R., and Axelrod, J.D. (2008). Asymmetric homotypic interactions of the atypical cadherin flamingo mediate intercellular polarity signaling. Cell 133, 1093–1105.

Chen, Z., Oh, D., Dubey, A.K., Yao, M., Yang, B., Groves, J.T., and Sheetz, M. (2018). EGFR family and Src family kinase interactions: mechanics matters? Curr Opin Cell Biol 51, 97–102.

Chuykin, I., Itoh, K., Kim, K., and Sokol, S.Y. (2021). Frizzled3 inhibits Vangl2-Prickle3 association to establish planar cell polarity in the vertebrate neural plate. J Cell Sci 134.

Courbard, J.R., Djiane, A., Wu, J., and Mlodzik, M. (2009). The apical/basal-polarity determinant Scribble cooperates with the PCP core factor Stbm/Vang and functions as one of its effectors. Dev Biol 333, 67–77.

Das, G., Jenny, A., Klein, T.J., Eaton, S., and Mlodzik, M. (2004). Diego interacts with Prickle and Strabismus/Van Gogh to localize planar cell polarity complexes. Development 131, 4467–4476.

Davey, C.F., and Moens, C.B. (2017). Planar cell polarity in moving cells: think globally, act locally. Development 144, 187–200.

Devenport, D. (2014). The cell biology of planar cell polarity. J Cell Biol 207, 171–179.

Devenport, D., and Fuchs, E. (2008). Planar polarization in embryonic epidermis orchestrates global asymmetric morphogenesis of hair follicles. Nat Cell Biol 10, 1257–1268.

Devenport, D., Oristian, D., Heller, E., and Fuchs, E. (2011). Mitotic internalization of planar cell polarity proteins preserves tissue polarity. Nat Cell Biol 13, 893–902.

Dobens, L.L., Shipman, A., and Axelrod, J.D. (2018). FijiWingsPolarity: An open source toolkit for semi-automated detection of cell polarity. Fly (Austin) 12, 23–33.

Espada, J., and Martin-Perez, J. (2017). An Update on Src Family of Nonreceptor Tyrosine Kinases Biology. Int Rev Cell Mol Biol 331, 83–122.

Gao, B., Song, H., Bishop, K., Elliot, G., Garrett, L., English, M.A., Andre, P., Robinson, J., Sood, R., Minami, Y., et al. (2011). Wnt signaling gradients establish planar cell polarity by inducing Vangl2 phosphorylation through Ror2. Dev Cell 20, 163–176.

Goodrich, L.V., and Strutt, D. (2011). Principles of planar polarity in animal development. Development 138, 1877–1892.

Guo, N., Hawkins, C., and Nathans, J. (2004). Frizzled6 controls hair patterning in mice. Proc Natl Acad Sci U S A 101, 9277–9281.

Guo, Y., Zanetti, G., and Schekman, R. (2013). A novel GTP-binding protein-adaptor protein complex responsible for export of Vangl2 from the trans Golgi network. eLife 2, e00160.

Humphries, A.C., and Mlodzik, M. (2018). From instruction to output: Wnt/PCP signaling in development and cancer. Curr Opin Cell Biol 51, 110–116.

Humphries, A.C., Narang, S., and Mlodzik, M. (2020). Mutations associated with human neural tube defects display disrupted planar cell polarity in Drosophila. eLife 9, e53532.

Jenny, A., Darken, R.S., Wilson, P.A., and Mlodzik, M. (2003). Prickle and Strabismus form a functional complex to generate a correct axis during planar cell polarity signaling. Embo J 22, 4409–4420.

Jenny, A., Reynolds-Kenneally, J., Das, G., Burnett, M., and Mlodzik, M. (2005). Diego and Prickle regulate Frizzled planar cell polarity signalling by competing for Dishevelled binding. Nat Cell Biol 7, 691–697.

Jessen, J.R., Topczewski, J., Bingham, S., Sepich, D.S., Marlow, F., Chandrasekhar, A., and Solnica-Krezel, L. (2002). Zebrafish trilobite identifies new roles for Strabismus in gastrulation and neuronal movements. Nat Cell Biol 4, 610–615.

Kelly, L.K., Wu, J., Yanfeng, W.A., and Mlodzik, M. (2016). Frizzled-Induced Van Gogh Phosphorylation by CK1epsilon Promotes Asymmetric Localization of Core PCP Factors in Drosophila. Cell reports 16, 344–356.

Koca, Y., Collu, G.M., and Mlodzik, M. (2022). Wnt-Frizzled planar cell polarity signaling in the regulation of cell motility. Curr Topics Dev Biol 150, in press.

Lei, Y.P., Zhang, T., Li, H., Wu, B.L., Jin, L., and Wang, H.Y. (2010). VANGL2 mutations in human cranial neural-tube defects. N Engl J Med 362, 2232–2235.

Lowell, C.A., Soriano, P., and Varmus, H.E. (1994). Functional overlap in the src gene family: inactivation of hck and fgr impairs natural immunity. Genes Dev 8, 387–398.

Marlow, F., Zwartkruis, F., Malicki, J., Neuhauss, S.C.F., Abbas, L., Weaver, M., Driever, W., and Solnica-Krezel, L. (1998). Functional interactions of genes mediating convergent extension, knypek and trilobite, during the partitioning of the eye primordium in zebrafish. Dev Biol 203, 382–393.

Montcouquiol, M., Rachel, R.A., Lanford, P.J., Copeland, N.G., Jenkins, N.A., and Kelley, M.W. (2003). Identification of Vangl2 and Scrb1 as planar polarity genes in mammals. Nature 423, 173–177.

Nikolopoulou, E., Galea, G.L., Rolo, A., Greene, N.D., and Copp, A.J. (2017). Neural tube closure: cellular, molecular and biomechanical mechanisms. Development 144, 552–566.

Ohata, S., Nakatani, J., Herranz-Perez, V., Cheng, J., Belinson, H., Inubushi, T., Snider, W.D., Garcia-Verdugo, J.M., Wynshaw-Boris, A., and Alvarez-Buylla, A. (2014). Loss of Dishevelleds disrupts planar polarity in ependymal motile cilia and results in hydrocephalus. Neuron 83, 558–571.

Park, M., and Moon, R.T. (2002). The planar cell polarity gene stbm regulates cell behaviour and cell fate in vertebrate embryos. Nat Cell Biol 4, 20–25.

Pedraza, L.G., Stewart, R.A., Li, D.M., and Xu, T. (2004). Drosophila Src-family kinases function with Csk to regulate cell proliferation and apoptosis. Oncogene 23, 4754–4762.

Ravni, A., Qu, Y., Goffinet, A.M., and Tissir, F. (2009). Planar cell polarity cadherin Celsr1 regulates skin hair patterning in the mouse. J Invest Dermatol 129, 2507–2509.

Seifert, J.R., and Mlodzik, M. (2007). Frizzled/PCP signalling: a conserved mechanism regulating cell polarity and directed motility. Nature reviews 8, 126–138.

Seo, H.S., Habas, R., Chang, C., and Wang, J. (2017). Bimodal regulation of Dishevelled function by Vangl2 during morphogenesis. Human molecular genetics 26, 2053–2061.

Shi, D., Komatsu, K., Hirao, M., Toyooka, Y., Koyama, H., Tissir, F., Goffinet, A.M., Uemura, T., and Fujimori, T. (2014). Celsr1 is required for the generation of polarity at multiple levels of the mouse oviduct. Development 141, 4558–4568.

Shinohara, K., and Hamada, H. (2017). Cilia in Left-Right Symmetry Breaking. Cold Spring Harbor perspectives in biology 9.

Simon, M.A., Kornberg, T.B., and Bishop, J.M. (1983). Three loci related to the src oncogene and tyrosine-specific protein kinase activity in Drosophila. Nature 302, 837–839.

Simons, M., and Mlodzik, M. (2008). Planar cell polarity signaling: from fly development to human disease. Annual review of genetics 42, 517–540.

Sirvent, A., Mevizou, R., Naim, D., Lafitte, M., and Roche, S. (2020). Src Family Tyrosine Kinases in Intestinal Homeostasis, Regeneration and Tumorigenesis. Cancers (Basel) 12.

Stahley, S.N., Basta, L.P., Sharan, R., and Devenport, D. (2021). Celsr1 adhesive interactions mediate the asymmetric organization of planar polarity complexes. eLife 10.

Stein, P.L., Vogel, H., and Soriano, P. (1994). Combined deficiencies of Src, Fyn, and Yes tyrosine kinases in mutant mice. Genes Dev 8, 1999–2007.

Strutt, D.I. (2001). Asymmetric localization of frizzled and the establishment of cell polarity in the Drosophila wing. Molecular Cell 7, 367–375.

Strutt, H., Gamage, J., and Strutt, D. (2019). Reciprocal action of Casein Kinase Iepsilon on core planar polarity proteins regulates clustering and asymmetric localisation. eLife 8.

Taylor, J., Abramova, N., Charlton, J., and Adler, P.N. (1998). Van Gogh: a new Drosophila tissue polarity gene. Genetics 150, 199–210.

Thomas, S.M., Soriano, P., and Imamoto, A. (1995). Specific and redundant roles of Src and Fyn in organizing the cytoskeleton. Nature 376, 267–271.

Tissir, F., Qu, Y., Montcouquiol, M., Zhou, L., Komatsu, K., Shi, D., Fujimori, T., Labeau, J., Tyteca, D., Courtoy, P., et al. (2010). Lack of cadherins Celsr2 and Celsr3 impairs ependymal ciliogenesis, leading to fatal hydrocephalus. Nat Neurosci 13, 700–707.

Torban, E., Wang, H.J., Groulx, N., and Gros, P. (2004). Independent mutations in mouse Vangl2 that cause neural tube defects in looptail mice impair interaction with members of the Dishevelled family. J Biol Chem 279, 52703–52713.

Vladar, E.K., Bayly, R.D., Sangoram, A.M., Scott, M.P., and Axelrod, J.D. (2012). Microtubules enable the planar cell polarity of airway cilia. Curr Biol 22, 2203–2212.

Wallingford, J.B. (2006). Planar cell polarity, ciliogenesis and neural tube defects. Human molecular genetics 15 Spec No 2, R227–234.

Wang, M., Marco, P., Capra, V., and Kibar, Z. (2019). Update on the Role of the Non-Canonical Wnt/Planar Cell Polarity Pathway in Neural Tube Defects. Cells 8.

Wang, Y., and Nathans, J. (2007). Tissue/planar cell polarity in vertebrates: new insights and new questions. Development 134, 647–658.

Wolff, T., and Rubin, G.M. (1998). strabismus, a novel gene that regulates tissue polarity and cell fate decisions in Drosophila. Development 125, 1149–1159.

Wu, J., and Mlodzik, M. (2008). The frizzled extracellular domain is a ligand for Van Gogh/Stbm during nonautonomous planar cell polarity signaling. Dev Cell 15, 462–469.

Wu, J., and Mlodzik, M. (2017). Wnt/PCP Instructions for Cilia in Left-Right Asymmetry. Dev Cell 40, 423–424.

Yanfeng, W.A., Berhane, H., Mola, M., Singh, J., Jenny, A., and Mlodzik, M. (2011). Functional dissection of phosphorylation of Disheveled in Drosophila. Dev Biol 360, 132–142.

Yang, Y., and Mlodzik, M. (2015). Wnt-Frizzled/planar cell polarity signaling: cellular orientation by facing the wind (Wnt). Annu Rev Cell Dev Biol 31, 623–646.

Ybot-Gonzalez, P., Savery, D., Gerrelli, D., Signore, M., Mitchell, C.E., Faux, C.H., Greene, N.D., and Copp, A.J. (2007). Convergent extension, planar-cell-polarity signalling and initiation of mouse neural tube closure. Development 134, 789–799.

