## Supplemental Figures for "A Van Gogh/Vangl tyrosine phosphorylation switch regulates its interaction with core Planar Cell Polarity factors Prickle and Dishevelled"

**Supplemental Information:**

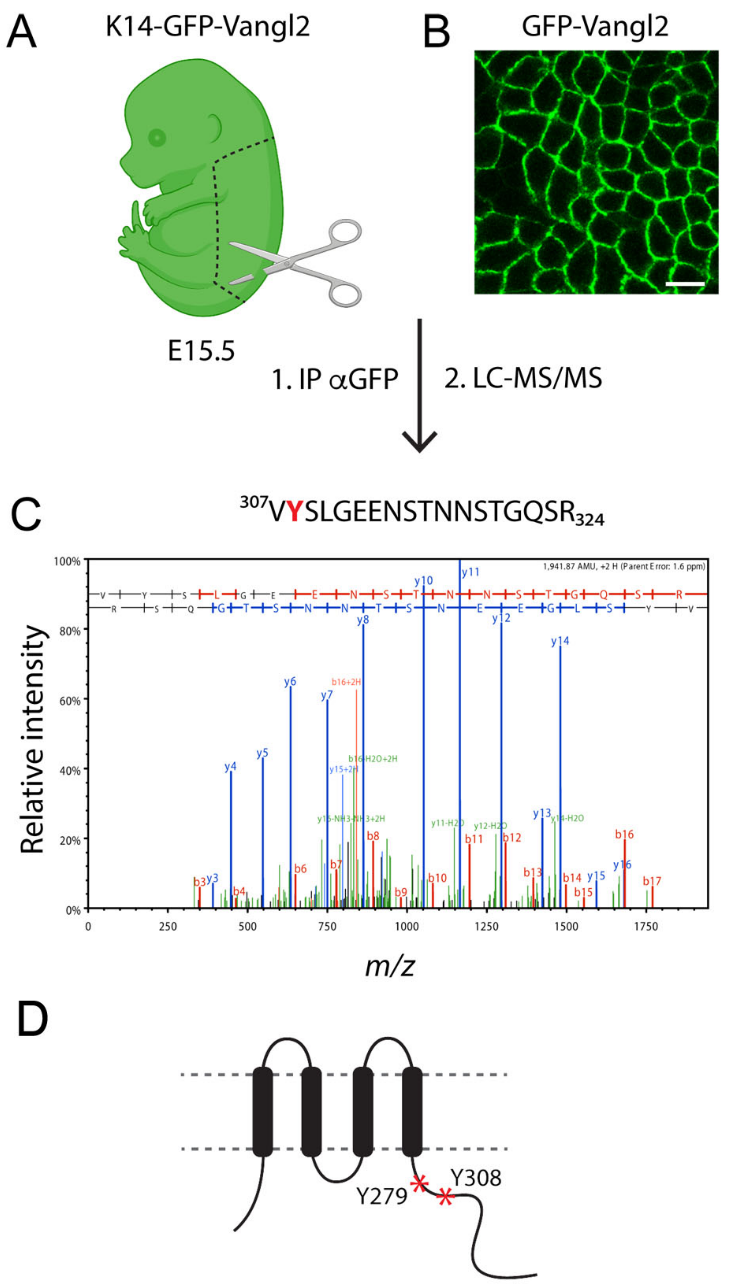


**Figure S1 (Supplement to Figure 1):**

**IP-MS approach from mouse skin to identify Vangl2 PTMs.**

**A**. Skins were dissected from E15.5 embryo expressing GFP-Vangl2 under the K14 skin-specific promoter. Epidermal lysates were prepared from frozen and cryo-milled skin samples and GFP-Vangl2 immunoprecipitated using anti-GFP antibodies. Created with BioRender.com

**B.** Planar view of basal layer from K14-GFP-Vangl2 whole mount epidermis. GFP-Vangl2 is localized to cell junctions. Scale bar 10um.

**C.** Representative m/z spectrum of Y308-contining peptide (aa307-394).

**D.** Schematic of mouse Vangl2 protein with asterisks indicating positions of phosphorylated tyrosines Y279 and Y308.

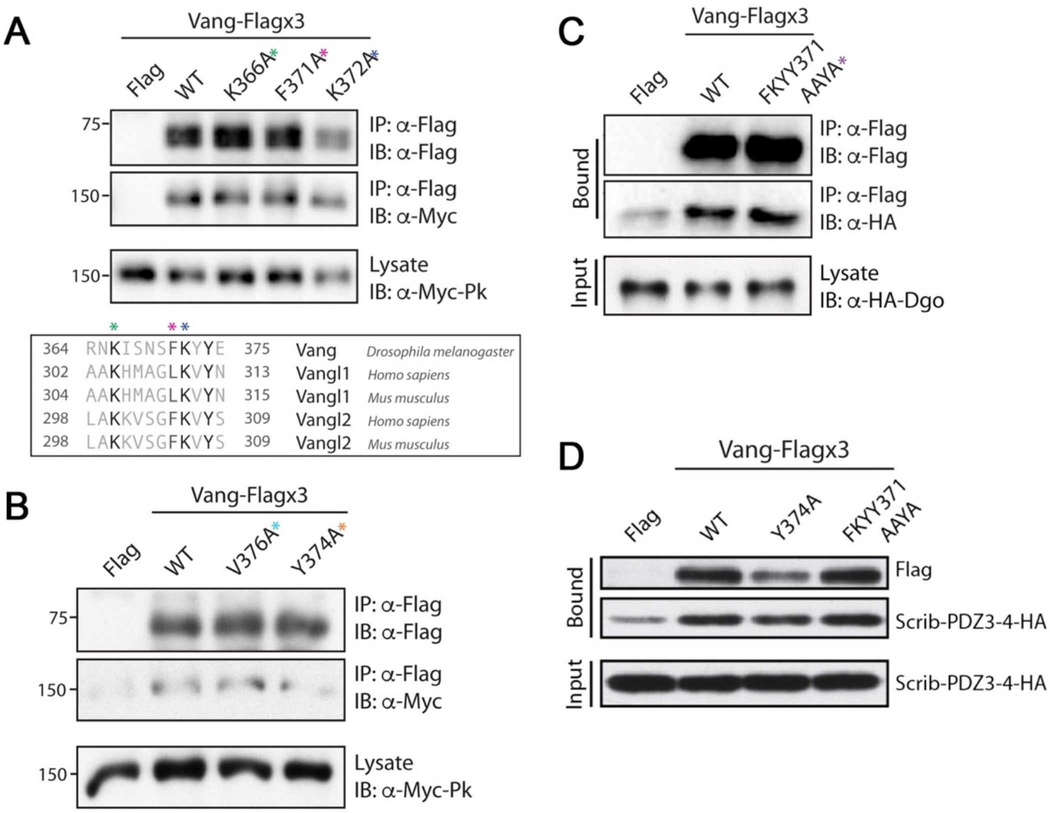


**Figure S2 (Supplement to Figure 2):**

**Specificity of Pk and Dsh binding to the Vang region 364-387**

To further define the binding site, we focused on the conservation within this region, as the mammalian orthologues, VANGL1/2 and PK family members also physically interact. A series of point mutations, substituting each conserved or partially conserved residue for alanine was generated and these experiments revealed that substitutions at K366, F371 and K372 had little effect alone: (**A**) Western blot showing binding between Myc-Pk and selected Vang-Flagx3 point mutants as indicated (see sequence alignment below, with colored asterisks highlighting the different amino acids mutated in the experiment). Note binding is retained in each case. (**B**) Western blot showing binding between Myc-Pk and Vang-Flagx3 V376A point mutant, which abrogates Vang-Dsh binding (see main Figure 2E for comparison; see main Figure 2F for sequence alignment with the colored corresponding asterisks highlighting the amino acids mutated). Note that Pk binding is retained in V376A case (cf to WT and Y374A as controls).

(**C**) and (**D**) Control binding experiments for Vang with additional PCP core factors/effectors, Dgo and Scribble (Scrib), demonstrating that neither Dgo nor Scrib bind to the region where Pk and Dsh interact. Note that mutants in the 371-FKYY-374 motif do not affect the Vang-Dgo and Vang-Scrib interactions.

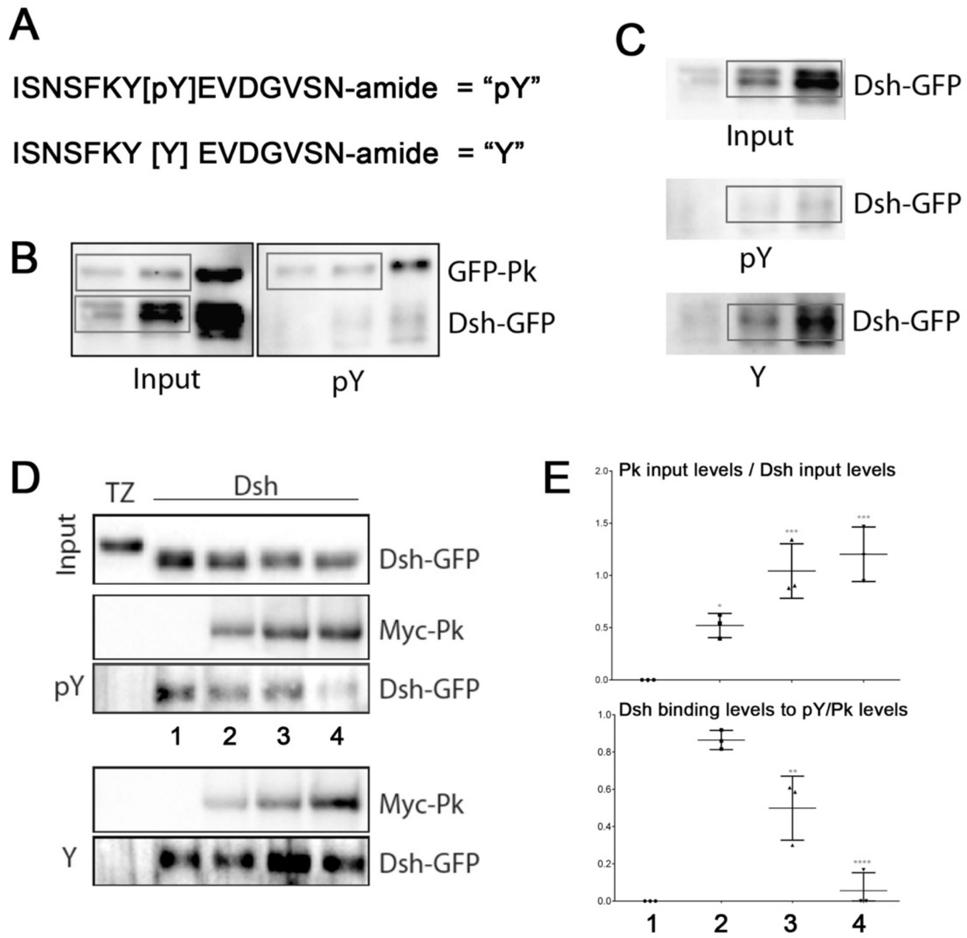


**Figure S3 (Supllement to Figure 3):**

**Y374 phospho-peptide binds preferentially to Pk.**

(**A**) Sequence of the synthetic peptides used [ISNSFKY[*pY*]EVDGVSN-amide] and the equivalent non-phosphorylated control, with the phospho-peptide denoted as “pY” and the control peptide as “Y” in the subsequent panels. (**B**) Pk interacts with/binds to the phosphorylated “pY” peptide significantly better than Dsh. (**C**) Dsh preferentially interacts with the non-phosphorylated “Y”-peptide.

(**D**) Binding of Dsh (Dsh-GFP is outcompeted by Pk (myc-Pk) on the “p-Y” (phospho-peptide), but not the non-phosphorylated (“Y”) peptide. Western blot showing retention of Dsh-GFP on beads coupled with the respective peptide, either “pY” for “Y”. Upper blot shows stable Dsh-GFP input, TZ is an unrelated control protein. Note reduced binding of Dsh-GFP when Pk levels are increased in the context of the pY peptide, but not the Y peptide. (**E**) Quantification of binding competition. Note that increasing PK levels causes a decrease in Dsh binding to the “pY” peptide.

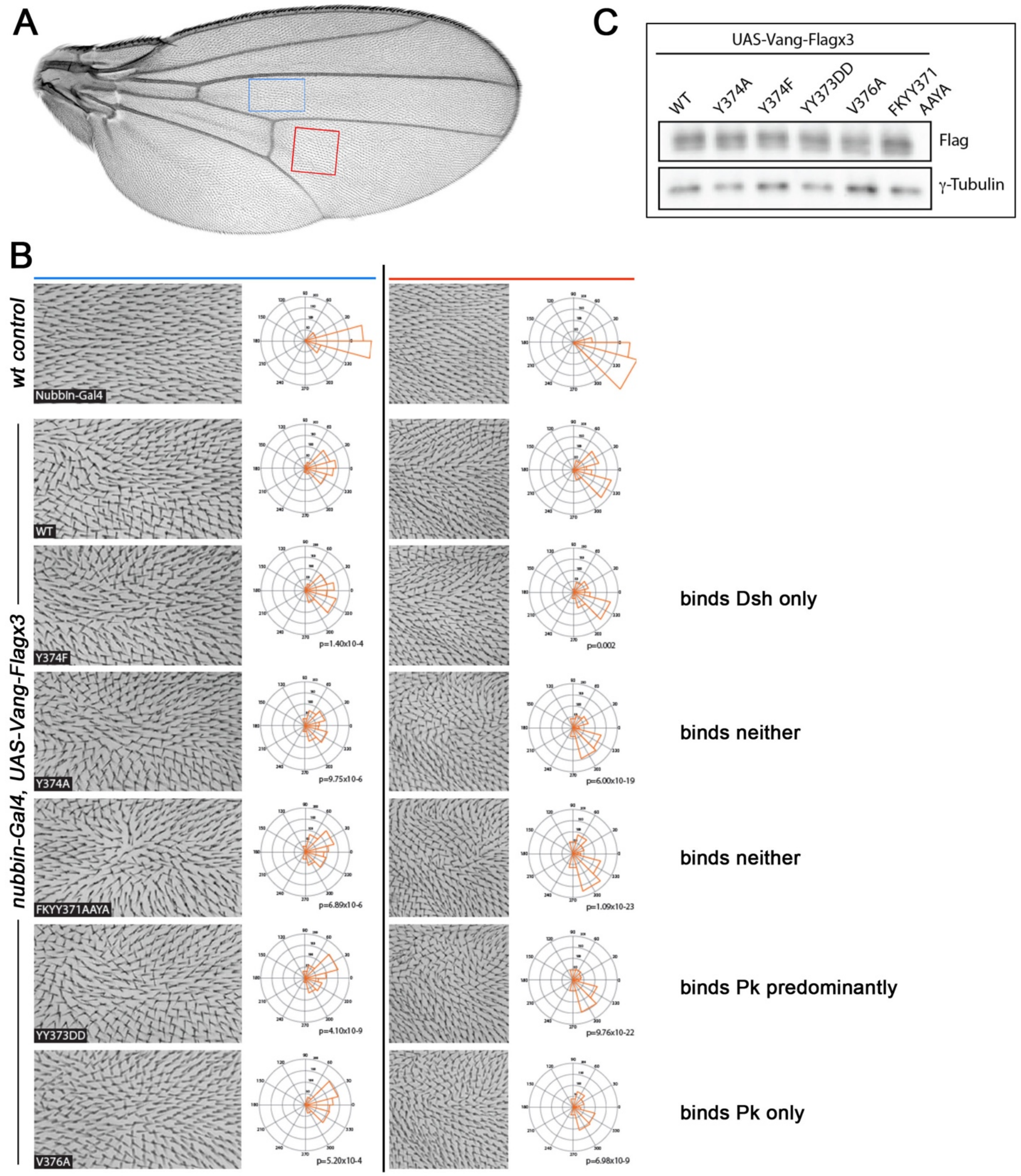


**Figure S4 (Supplement to Figure 4):**

**Distinct *in vivo* gain-of-function behavior of Vang and the respective binding mutants.**

Different behavior of the individual Vang binding mutants *in vivo*, as assayed in a GOF scenario in developing wings. (**A**) wild-type wing overview picture with boxed areas of regions of interest (ROI) in blue box and red box, as shown in (**B**) at higher magnification. In panels (**B**), each ROI (blue on left and red on right) was analyzed for cellular polarity by actin hair orientation (genotypes are as indicated, all transgenes were expressed under the control of *nubbin-Gal4*). Quantification of polarity angle distribution is shown on right of representative panel for each genotype. 3-6 animals were analyzed per genotype. Note that each different mutant shows a specific, but distinct distribution (as compared to Vang-*WT* and other mutants; *p* values vs Vang-*WT* are shown, as determined by Chi-squared tests). For example, note different polarity distributions in Vang-Y374F (binding Dsh only) to Vang-YY373DD (binding predominantly Pk) or Vang-Y374A (binding neither Pk nor Dsh), with the exception of Vang-YY373DD and Vang-V376A, which look similar and also both bind Pk only. (**C**) Western blot of wings from the indicated genotypes, demonstrating highly comparable and even expression levels of all Vang isoforms (wild-type and mutant).
